# Co-zorbs: Motile, multispecies biofilms aid transport of diverse bacterial species

**DOI:** 10.1101/2024.08.29.607786

**Authors:** Shruthi Magesh, Jonathan H. Schrope, Nayanna Mercado Soto, Chao Li, Amanda I. Hurley, Anna Huttenlocher, David J. Beebe, Jo Handelsman

**Affiliations:** Wisconsin Institute for Discovery and Department of Plant Pathology, University of Wisconsin-Madison; Madison, WI, USA; Microbiology Doctoral Training Program, University of Wisconsin-Madison; Madison, WI, USA; Department of Biomedical Engineering, University of Wisconsin-Madison; Madison, WI, USA; Department of Pathology and Laboratory Medicine, University of Wisconsin-Madison; Madison, WI, USA; Department of Medical Microbiology and Immunology, University of Wisconsin-Madison; Madison, WI, USA; Carbone Cancer Center, University of Wisconsin-Madison; Madison, WI, USA; Avantiqor, 800 Wharf St SW, Washington, DC 20024

**Keywords:** *Flavobacterium johnsoniae*, polymicrobial biofilms, tri-zorb, spatial organization, zebrafish

## Abstract

Biofilms are three-dimensional structures containing one or more bacterial species embedded in extracellular polymeric substances. Although most biofilms are stationary, *Flavobacterium johnsoniae* forms a motile spherical biofilm called a zorb, which is propelled by its base cells and contains a polysaccharide core. Here, we report formation of spatially organized, motile, multispecies biofilms, designated “co-zorbs,” that are distinguished by a core-shell structure. *F. johnsoniae* forms zorbs whose cells collect other bacterial species and transport them to the zorb core, forming a co-zorb. Live imaging revealed that co-zorbs also form in zebrafish, thereby demonstrating a new type of bacterial movement in vivo. This discovery opens new avenues for understanding community behaviors, the role of biofilms in bulk bacterial transport, and collective strategies for microbial success in various environments.

**Significance Statement:** This paper reports the discovery of co-zorbs, which are spherical aggregates of bacteria that move and transport other bacteria. Zorbs move toward other bacteria and collect them in a manner reminiscent of phagocytes. Once inside the zorb, the new species form a striking, organized core. The discovery of co-zorbs introduces an entirely new type of bacterial movement and transport involving cooperation among bacterial species. Co-zorbs have potential for engineering microbial systems for biotechnology applications and for managing spread of bacterial pathogens in their hosts.

## Introduction

Like a bustling city where a variety of people live together, interact with one another, and shape the environment, diverse microorganisms converge to create intricate structures called biofilms. Within these multispecies biofilms, microbes embedded in an extracellular matrix interact, influencing the spatial architecture and functionality of the community (1). These interactions give rise to emergent properties that are absent in single-species biofilms, profoundly impacting the overall behavior and characteristics of the biofilm (2). The significance of multispecies biofilms across applications in health care, agriculture, and environmental management highlights the need to understand the complex interspecies interactions in biofilms, which is crucial to address global challenges as diverse as food poisoning, biofouling in industrial infrastructure, and antibiotic resistance in chronic infections (3).

Biofilms in the environment are typically stationary structures. We recently discovered that *Flavobacterium johnsoniae*, a Gram-negative bacterium commonly found in soil and freshwater, uses cell features involved in gliding motility and colonization to form spherical biofilm-like microcolonies called zorbs (4, 5). These zorbs are unique as they are propelled by their base cells without pili or flagella and contain an extracellular polysaccharide core. There are only a few examples in which motility is observed in interspecies biofilms. For example, *Pseudomonas aeruginosa* shifts from collective to single-cell motility, initiating exploratory movements upon sensing *Staphylococcus aureus* biofilms (6), *Capnocytophaga gingivalis* transports non-motile bacteria as cargo along its cell length, shaping the spatial organization of polymicrobial communities (7), and *Candida albicans* and *Streptococcus mutans* demonstrate a “forward-leaping motion” in interkingdom biofilms (8). These behaviors depend on either single-cell motility, hitchhiking, or interspecies expansion, in which individual cells carrying non-motile bacteria move, grow, and spread across surfaces to form interspecies or multispecies biofilms.

In contrast, our study presents a new mechanism of bacterial movement in which motile *F. johnsoniae* biofilms transport both motile and non-motile bacterial species in a core-shell structure. In this unique collective movement, a group of *F. johnsoniae* cells surround and collect other bacterial species and transports them as a multispecies biofilm structure across surfaces, including inside zebrafish larvae. Co-zorbing challenges the notion that multispecies biofilms are stationary and suggests a fertile area for future research in community ecology and infectious disease.

## Results

### Structure and formation of spatially organized motile multispecies biofilms

To study the interaction of *F. johnsoniae* zorbs with other bacterial species, we introduced *F. johnsoniae* and *E. coli* into an under-oil microfluidic device (4-5, 9–11) amenable to long term timelapse imaging of small-volume cultures (2 µL for >18 hours) without evaporation. In monoculture, *F. johnsoniae* formed zorbs and *E. coli* did not. When the two species were co-inoculated, they interacted and formed a structure we designate as a multispecies “co-zorb” in which the *E. coli* occupied the core surrounded by a shell of *F. johnsoniae* (Fig. 1*A* and *B*). In a 17-hour study of co-zorb growth, we found that smaller co-zorbs with a maximum diameter of approximately 27 +/- 6 microns appeared at ∼6 hours and merged into larger co-zorbs by 12-15 hours, reaching a maximum diameter of approximately 105 +/- 18 microns before dispersing around 16 hours (Movie S1). Confocal imaging of co-zorbs revealed a distinctive three-dimensional core-shell structure formed by the two species (Fig. 1*B*). Notably, co-zorbs of *F. johnsoniae* and *E. coli* were motile (Fig. 1*C*) and traversed greater distances than monospecies zorbs (Fig. 1*D*), suggesting that the presence of *E. coli* enhances the collective movement of the biofilm structure.

**Figure 1.**
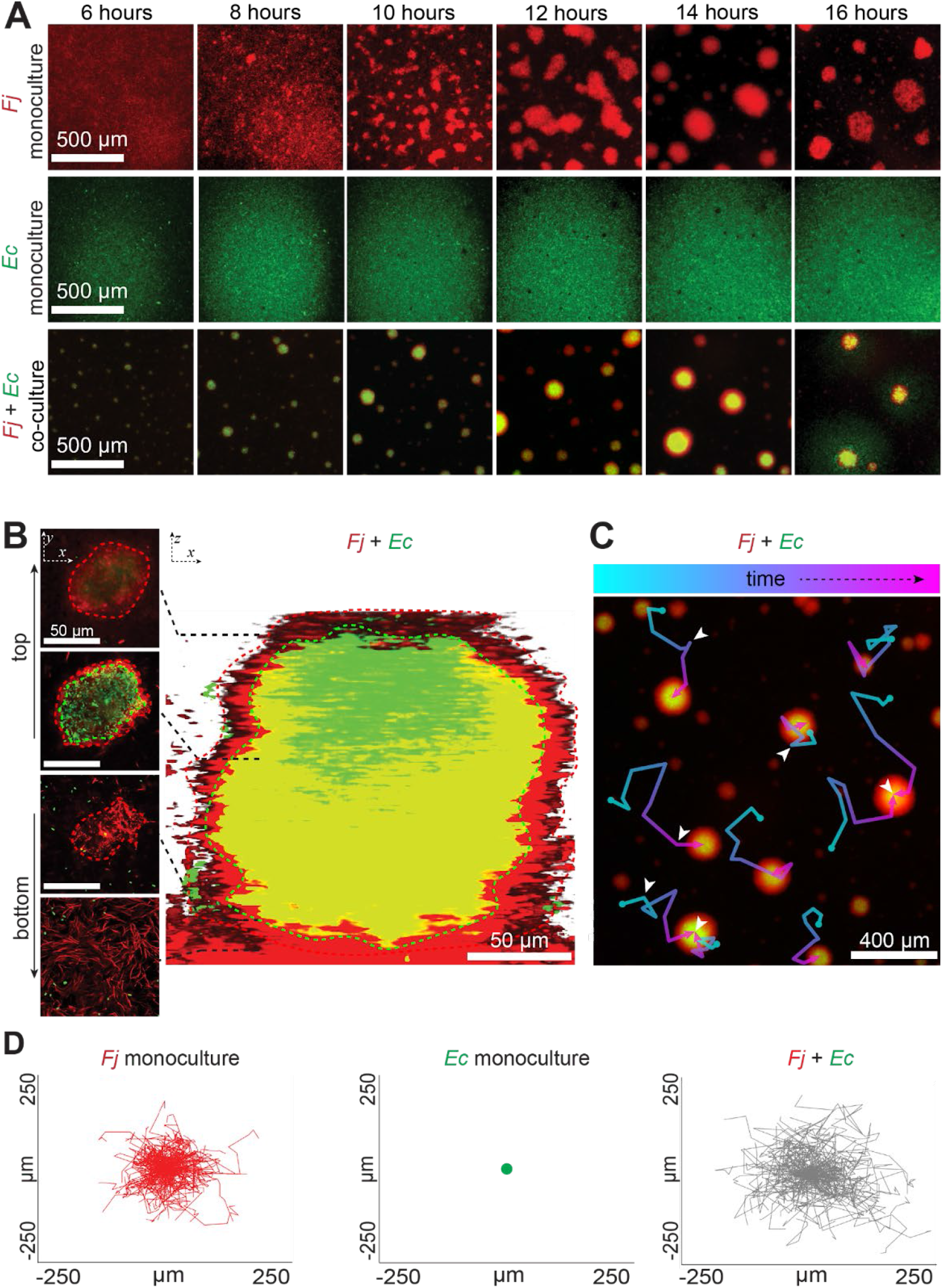
Structure and formation of co-zorbs. (A) Formation of co-zorbs. Timelapse images showing formation of *F. johnsoniae (Fj)* zorbs (red), *E. coli (Ec)* (green) and *F. johnsoniae*-*E. coli (Fj+Ec)* co-zorb (red-green) over 16 hours. (B) *E. coli* is localized within the core of the *F. johnsoniae* zorb. Confocal reconstruction showing the core-shell structure of mature co-zorb (right) with corresponding two-dimensional images (left) at various z-positions along the height of the co-zorb. (C) Co-zorbs are motile. Tracks of co-zorbs over time, with merging events (when two co-zorbs combine into one) denoted by a white arrow. (D) Co-zorbs traverse a greater distance than zorbs. Plots of track displacement for *F. johnsoniae* zorbs, *E. coli*, and co-zorbs from 6 to 17 hours with the origin (i.e., x = y = 0 μm) as the starting point of zorb movement. Red represents the tracks of *F. johnsoniae* zorbs, green represents tracks of *E. coli*, and gray represents displacement tracks of co-zorbs (*F. johnsoniae* + *E. coli*).

### Several bacterial species form co-zorbs with *F. johnsoniae*

To determine whether *F. johnsoniae* forms co-zorbs with other bacterial species, we tested fluorescently labeled strains of several species. These included four Gram-positive members of the Bacillota phylum (*Bacillus cereus, Bacillus subtilis, Listeria monocytogenes*, and *Staphylococcus aureus*) and three Gram-negative members of the Pseudomonadota in addition to *E. coli* (*Agrobacterium tumefaciens, Salmonella enterica*, and *Pseudomonas aeruginosa*). These strains represent bacteria with diverse characteristics: motile, non-motile, sporulating, non-sporulating, pathogenic, and non-pathogenic. Remarkably, despite their diverse characteristics, all bacterial species tested with *F. johnsoniae* formed motile co-zorbs (Fig. 2*A*) with variations in core area, core centrality (normalized distance to zorb center), co-zorb circularity, and speed (Fig. 2*B* and Fig. S1), suggesting the significant influence of interspecies interactions in the nature of co-zorbs.

**Figure 2.**
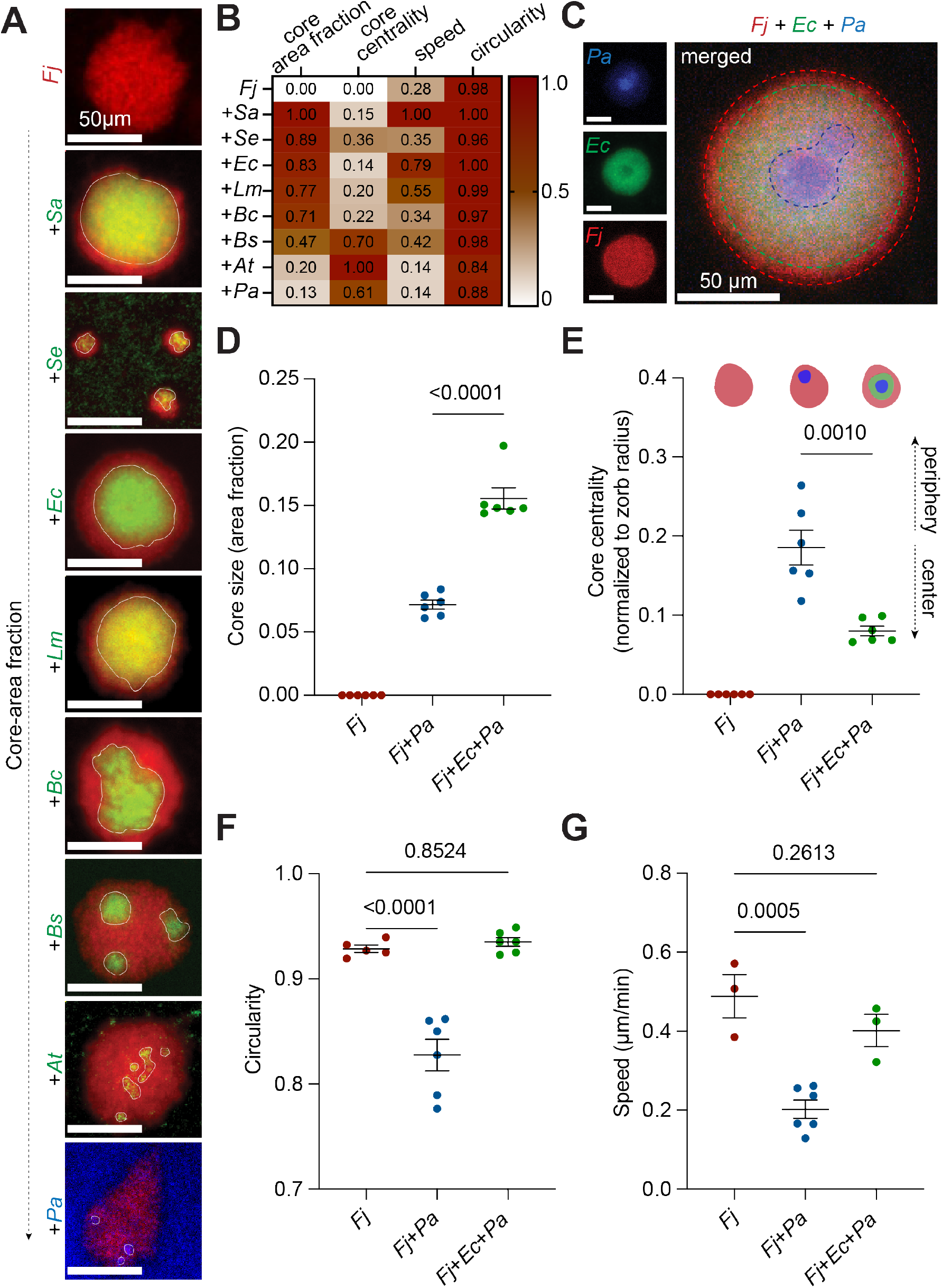
Multi-species zorbing dynamics: Co-zorbing and tri-zorbing with *F. johnsoniae* (A) Co-zorbs with *F. johnsoniae* (*Fj*) in red and second species in green or blue; arranged in order of core-area fraction (area occupied by second species divided by total co-zorb area). Top to bottom: *Fj* zorbs, *Fj* co-zorbs with *S. aureus* (*Sa*), *S. enterica* (*Se*), *E. coli* (*Ec*), *L. monocytogenes* (*Lm*), *B. cereus* (*Bc*), *B. subtilis* (*Bs*), *A. tumefaciens* (*At*), and *P. aeruginosa* (*Pa*). (B) Values represent three biological replicates normalized to the maximum mean value in each category. (C) Tri-zorbing with *Pa* and *Ec. Fj* (red, shell), *E. coli* (green, outer-core), and *P. aeruginosa* (blue, inner-core). (D) *Pa* occupies greater core area in tri-zorbs than in co-zorbs. y-axis indicates core area of *Pa* (core area/total area). (E) Distance between core and center of zorb normalized to radius of zorb. (F) Tri-zorbs are more circular (4π X area/perimeter^2^) than *Fj-Pa* co-zorbs. (G) Tri-zorbs move faster than *Fj-Pa* co-zorbs. Mean zorb, co-zorb, and tri-zorb speed (µm/min). In D-G, each data point represents a biological replicate (n=10 measurements per replicate). Error bars indicate S.E.M. For circularity and speed, significance is determined by one-way ANOVA with comparisons to *Fj* alone. For core size and distance from center, significance determined by unpaired Student’s t-test assuming normal distributions.

We then sought to determine whether *F. johnsoniae* exhibited a species preference for co-zorbing. Hence, we co-inoculated *F. johnsoniae* with *E. coli* and *P. aeruginosa* and monitored their dynamics for over 18 hours (**<u>Movie 1</u>**). Instead of preferentially co-zorbing with one species, co-inoculation of all three species resulted in the formation of tri-zorbs, with *E. coli* and *P. aeruginosa* arranged in a concentric pattern within the *F. johnsoniae* zorb (Fig. 2*C*). In the presence of *E. coli, P. aeruginosa* occupied more area within the tri-zorb than in the *F. johnsoniae*-*P. aeruginosa* co-zorb (Fig. 2*D*) and was localized closer to the center (Fig. 2*E*), suggesting that *E. coli* influences the spatial organization of *P. aeruginosa* in tri-zorbs. In addition, tri-zorbs were more circular (Fig. 2*F*) and moved faster than the *F. johnsoniae*-*P. aeruginosa* co-zorbs and at a speed comparable to *F. johnsoniae* alone (Fig. 2*G*). Together, these data suggest that the presence of *E. coli* influences the interaction between *F. johnsoniae* and *P. aeruginosa*, thereby affecting the overall dynamics of the biofilm structure and behavior. Thus, tri-zorbing demonstrates an emergent property of the community since this higher order interaction could not be predicted from the individual members. Additionally, we observed tri-zorbs when *F. johnsoniae* and *P. aeruginosa* were co-inoculated with the Gram-positive, non-motile *S. aureus*, in which *P. aeruginosa* again localized in the center, suggesting that the spatial organization within tri-zorbs is not random (Fig. S2).

### *F. johnsoniae* aggregates, localizes, and transports the second species

To determine which species drives co-zorb formation, we used staggered inoculation to allow zorbs to form before introducing the second species. *F. johnsoniae* was incubated for 8-10 hours until zorbs formed, and then either *E. coli* (Gram-negative and motile) or *S. aureus* (Gram-positive and non-motile) was added. Time-lapse imaging showed that both *E. coli* and *S. aureus* localized within the core of *F. johnsoniae* zorbs when added to pre-formed zorbs (Fig. 3*A*). This observation led us to formulate two hypotheses regarding the mechanism underlying co-zorbing: either (1) the second species penetrates the *F. johnsoniae* zorb structure or (2) *F. johnsoniae* recruits and localizes the second species within its core. To distinguish between these hypotheses, we imaged zorbs at a high magnification at 15-second intervals (Fig. 3*B*). Upon inoculation with either *E. coli* or *S. aureus, F. johnsoniae* cells surrounded cells of the second species, forming a small aggregate of approximately 5-10 cells around it. Several of these small aggregates carrying the second species merged with other aggregates either carrying a second species or not, ultimately leading to the formation of co-zorbs with distinct core-shell structures (**<u>Movie 2</u>** and Movie S2). These observations indicate that *F. johnsoniae* zorbs recruit other species to the zorb, thereby supporting the second hypothesis (Fig. 3*C*).

**Figure 3.**
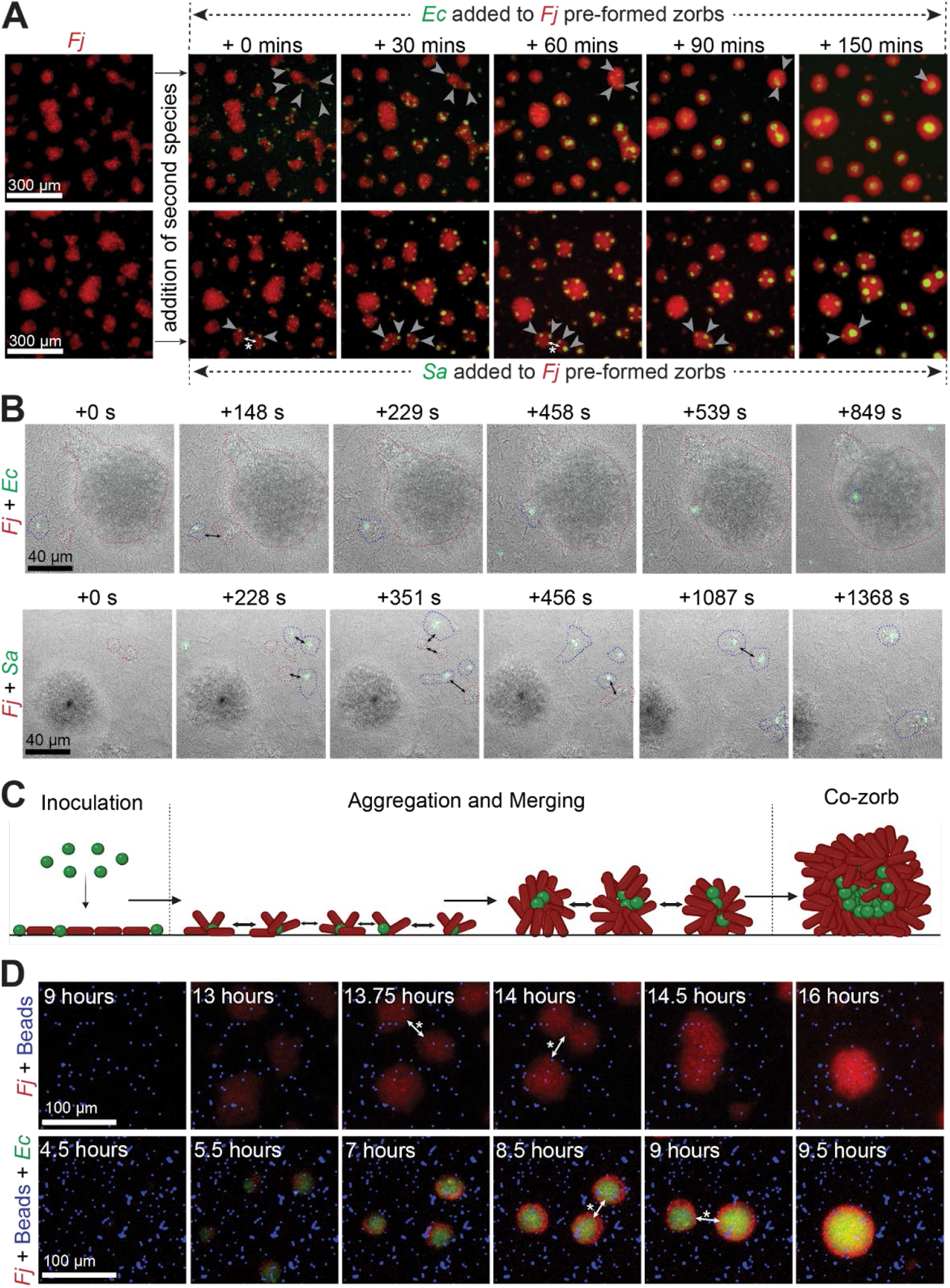
*F. johnsoniae* aggregates, localizes, and transports the second species (A) Pre-formed *F. johnsoniae* zorbs co-zorb with *E. coli* and *S. aureus*. Timelapse images showing co-zorbing after staggered inoculation of *E. coli* (top) or *S. aureus* (bottom) with pre-formed *F. johnsoniae* zorbs. (B) *F. johnsoniae* aggregates around the second bacterial species and localizes it within the zorb core. Timelapse images showing that *F. johnsoniae* cells (bright-field) on the surface of the plate aggregate and transport *E. coli* (top) and *S. aureus* (bottom) (green). Blue circles highlight *F. johnsoniae* cells surrounding the second species; the red circles highlight *F. johnsoniae* aggregates merging with them. Merging events between single or multi-species aggregates are denoted by a double arrow. (C) Schematic of the process of co-zorbing (Created with Biorender.com). Left to right: After adding the second species (green), *F. johnsoniae* cells (red) on the bottom of the plate aggregate around the second species, and move the second species from one point to another. These aggregates merge, forming larger aggregates, which eventually develop into co-zorbs with distinctive core-shell structure. (D) *F. johnsoniae* does not co-zorb with polystyrene beads. Timelapse image showing *F. johnsoniae* (red) preferentially co-zorbing with *E. coli* (green) and not polystyrene beads (blue).

To determine whether motility of either partner is required for co-zorb formation, we tested non-motile mutants. A non-motile *F. johnsoniae* Δ*sprA* mutant did not form zorbs alone and did not form co-zorbs with either *E. coli* or *S. aureus* (Fig. S3*A*). The cells of the non-motile mutant failed to collect and transport the second species, highlighting the importance of *F. johnsoniae* motility for co-zorbing and further supporting the model that *F. johnsoniae* actively recruits and localizes the second species within the zorb core.

In contrast to *F. johnsoniae*, motility of the second species is not required for co-zorbing. *E. coli* mutants containing deletions in genes required for motility, *fliC* and *fliD*, or chemotaxis, *cheB* and *cheZ*, co-zorbed with *F. johnsoniae* (Fig. S3*B*). Combined with the finding that non-motile species such as *S. aureus* form co-zorbs, this result suggests that the motility of the second species is not required for co-zorbing. Additionally, *F. johnsoniae* did not co-zorb with fluorescent polystyrene beads when they were added either alone or in the presence of *E. coli* (Fig. 3*D*), suggesting that recognition or specificity may play a role. Together these results indicate that *F. johnsoniae* is the active partner in co-zorbing—it must be motile, and it drives rapid translocation—whereas the second species can co-zorb whether it is motile or non-motile, suggesting it is the passive partner in the process.

### Co-zorbing enhances bacterial transport in zebrafish

We sought to determine whether co-zorbs form and move in an animal host. Due to its optical transparency and thus amenability to live fluorescence imaging, the larval zebrafish has emerged as a powerful model system to visualize host-microbe interactions (12,13). Moreover, *F. johnsoniae* has been implicated in fish diseases (14) and infects zebrafish (15), making it an ideal system in which to study zorbs. We injected bacteria into the hindbrains of larval zebrafish two days after fertilization (2dpf) (16,17) (Fig. 4*A*) and observed the formation of distinct core-shell structures between *F. johnsoniae* and either *E. coli* or *S. aureus* (Fig. 4*B*, Fig. 4*C* and Movie S3). The co-zorbs were motile, thereby appearing to recapitulate the in vitro behavior (Fig. 4*D*). Moreover, in the zebrafish larvae, *E. coli* and *S. aureus* that co-zorbed with *F. johnsoniae* moved faster than their monoculture counterparts (*E. coli* or *S. aureus* alone) (Fig. 4*E*). The ability of *F. johnsoniae* to form motile multispecies biofilms and maintain spatial organization within the zebrafish hindbrain indicates that this phenomenon can be extended to complex biological environments and warrants further investigation.

**Figure 4.**
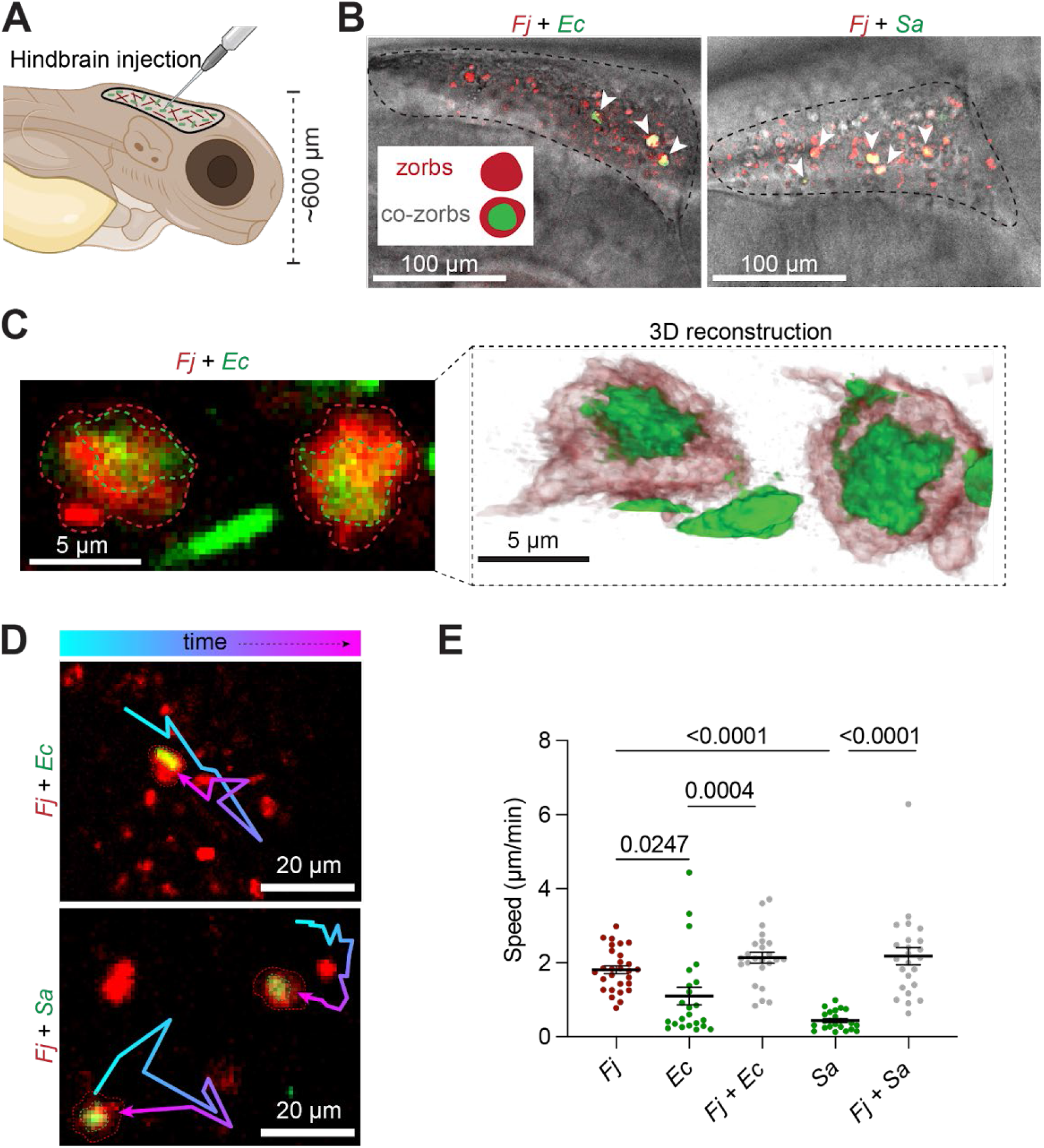
Co-zorbs retain spatial organization and motility in a live animal. (A) Schematic of larval zebrafish hindbrain injection. *F. johnsoniae* (10^9^ CFU/mL) and *E. coli* (10^9^ CFU/mL) or *F. johnsoniae* (10^9^ CFU/mL) and *S. aureus* (10^9^ CFU/mL) were injected into the hindbrain of two days post fertilization (2dpf) larval zebrafish. (B) Formation of co-zorbs inside zebrafish hindbrain. (Left) *F. johnsoniae-S. aureus* co-zorbs; (Right) *F. johnsoniae*-*E. coli* co-zorbs. (C) Confocal reconstruction shows co-zorbs form core-shell structure in vivo. (Left) Confocal image of *F. johnsoniae*-*E*.*coli* co-zorb injected at cell-density of 10^10^ CFU/mL. (Right) 3D reconstruction of *F. johnsoniae*-*E*.*coli* co-zorb observed in vivo. (D) Co-zorbs are motile within zebrafish. An image showing tracks of (top) *F. johnsoniae*-*E*.*coli* co-zorbs and (bottom) *F. johnsoniae-S. aureus* co-zorbs over 48 min. (E) Co-zorbs enable transport of *E. coli* or *S. aureus* at significantly greater speed than their monoculture counterparts (*E. coli* or *S. aureus* alone). Each data point represents the mean speed of all cells or co-zorbs within the field of view for a single fish (n = 9 fish over three independent experiments); error bars denote mean +/- S.E.M. Significance was determined by one-way ANOVA with multiple comparisons between each experimental group.

## Discussion

In this study, we introduce a new bacterial behavior, designated co-zorbing, in which *F. johnsoniae* zorbs, or motile biofilms, encapsulate diverse bacterial species to enable collective transport in a spatially organized structure. This process illustrates cooperative motility of a biofilm structure, interspecies interactions, and spatial organization, providing new insights into the complexity of interspecies interactions among bacteria. The ability of *F. johnsoniae* to form co-zorbs with both Gram-positive and Gram-negative bacteria, despite their diverse characteristics, and to form tri-zorbs when co-inoculated with two companion species underscores the versatility and variety of zorb interactions. The formation of co-zorbs inside zebrafish indicates their potential for mediating biological interactions with a host.

Collective behavior is observed throughout the biological world. Birds flock, fish school, and ants form colonies, and at the microscopic level, animal cells coordinate movement during morphogenesis, tissue remodeling, and cancer progression (18,19). In prokaryotes, collective behavior is often associated with surfaces and involves diverse forms of motility, including swarming, twitching, gliding, and swimming (20). *F. johnsoniae* is a gliding bacterium that uses the Type IX Secretion System (T9SS) and gliding motility apparatus to move across surfaces, exhibiting collective motility in the formation of vortex patterns under low-nutrient conditions (21,22). Other members of the phylum Bacteroidota, such as *C. gingivalis*, swarm in counterclockwise vortex patterns and exhibit cargo transport of non-motile species within the oral microbiome (7). Members of the phylum *Myxococcota* coordinate swarming for predation and aggregate into multicellular fruiting bodies under nutrient-deprived conditions (23). Zorbs and co-zorbs add to the pantheon of group strategies that bacteria employ to navigate their environments.

Motility and biofilm formation are typically used by bacteria to adapt to very different environmental conditions, and consequently, there are few examples of motile biofilms, such as zorbs (24-26). We demonstrate the movement of single-species and multi-species biofilms, as well as the bulk localization and transport of diverse species by *F. johnsoniae*. Several collective behaviors of *F. johnsoniae* are inhibited by high glucose levels (22, 27–29), including zorb formation (Fig. S4), suggesting that such collective motility may be a strategic response to nutrient deprivation. For example, colony spreading in *F. johnsoniae* is inhibited by N-acetylglucosamine, a structural component of the bacterial cell wall peptidoglycan (22), which serves as a carbon source for *F. johnsoniae* under low nutrient conditions (30). Perhaps *F. johnsoniae*, well-known for its ability to degrade complex polymers (31, 32), uses co-zorbing as a nutrient-acquisition strategy, feeding on cell components of the encapsulated species while providing it with rapid, safe transit across large distances inside the zorb. The benefits of forming co-zorbs remain to be determined. In particular, the impact of co-zorbs on bacterial spread and colonization in animals, on plant surfaces, or in other environments is a fertile area for future investigation.

The genetic tractability of *F. johnsoniae* makes it a candidate for engineering for practical use. It might serve as a tool to encapsulate other bacteria, including *S. aureus*-MRSA, and act as a scavenger in a manner analogous to that of phagocytes. Co-zorbing therefore presents a new means of bacterial translocation, extends our knowledge of microbial community behaviors, and opens innovative avenues for biotechnological and medical applications.

## Materials and Methods

### Bacterial strains and growth conditions

Strains and growth conditions used in this study are listed in Table 1. Each strain was grown overnight in a shaking incubator under the conditions specified in Table 1.

**Table 1.** List of bacterial strains and plasmids used in this study.

| Species | Strain background | Plasmid | Description | Growth conditions | Source |
| --- | --- | --- | --- | --- | --- |
| *Flavobacterium johnsoniae* | UW101 | None | Wild type | 0.5X TSB at 28°C | (31) |
| *Flavobacterium johnsoniae*-FJmS101 | UW101 | mStrawberry gene expression plasmid pSCH443; Em<sup>r</sup> | Wild type | 0.5X TSB supplemented with 100 µg mL<sup>-1</sup> erythromycin at 28°C | Plasmid provided by Dr. E.D. Walker (33) |
| *Flavobacterium johnsoniae*-CJ2302ΔsprA | CJ1827<br>*rpsL2ΔsprA* (Sm<sup>r</sup>) | None | Mutant deficient in SprA, resulting in the inability to attach to surfaces and move | 1/2-strength TSB at 28°C | Provided by Dr. Mark McBride (34) |
| *E. coli* K-12-sJMP3053 | BW25141 | Super folded GFP gene expression plasmid pJMP2774; Amp<sup>r</sup> | Wild-type strain used for all experiments unless stated otherwise. sJMP3053 is a cloning strain with anticrispr in Tn7att site | LB supplemented with 100 µg mL<sup>-1</sup> ampicillin at 37°C | Provided by Dr. Jason Peters (35) |
| *Staphylococcus aureus*-MRSA | USA300-LAC | super folded highly expressed GFP gene expression plasmid pCM29; Cm<sup>r</sup> | Wild type | 0.5X TSB at 37°C | Provided by Dr. J.D. Sauer (36) |
| *Bacillus subtilis* | JDW 4245 | None | Wild type used for co-zorbing experiments. Contains genome integration: *amy E::PilvBbp4ΔT-GFPmut2* | 0.5X TSB at 28°C | Provided by Dr. Ju Wang |
| *Bacillus cereus* | UW-85 | GFP gene expression plasmid pAD123_31-26; Cm<sup>r</sup> | Wild type used for co-zorbing experiments | 0.5X TSB supplemented with 10 µg mL<sup>-1</sup> chloramphenicol at 28°C | (37) |
| *Listeria monocytogenes* | 10403s | None | Wild type strain used for co-zorbing experiments. It has a genome integration of pPL2 (GFP) | 0.5X TSB at 28°C | Provided by Dr. J.D. Sauer (38) |
| *Salmonella enterica* | sv. Typhimurium 14028s | GFP gene expression plasmid pKT-Kan, Kan<sup>r</sup> | Wild type strain used for co-zorbing experiments | 0.5X TSB supplemented with 25 µg mL<sup>-1</sup> kanamycin at 37°C | Provided by Dr. Jeri Barak (39) |
| *Agrobacterium tumefaciens* | C58 | GFP gene expression plasmid pJZ383, P<sub>tac</sub>::*gfpmut3*, Sp<sup>r</sup> | Wild type strain used for co-zorbing experiments | 0.5X TSB supplemented with 150 µg mL<sup>-1</sup> chloramphenicol at 28°C | Provided by Dr. Clay Fuqua (40) |
| *Pseudomonas aeruginosa*-PA84 CFP7 | PA01 | None | Wild type strain used for all experiments. It has a genomic integration of ECFP with Pa1/04/03 promoter in Tn7att site, Kan<sup>r</sup> | LB supplemented with 50 µg mL<sup>-1</sup> kanamycin at 28°C | From the Handelsman lab strain collection |
| *E. coli* K-12- sJMP2686 | BW25113 | None | Wild type strain for mutants in keio collection. It has a genomic integration of GFP (Pveg promoter) between *yjaA* and *yjaB*, *catS*, *kanS* sites | LB at 37°C | Provided by Dr. Jason Peters (41) |
| *E. coli* K-12-Δ*fliC*-45 | BW25113 | GFP gene expression plasmid pGEN-GFP(LVA); Amp<sup>r</sup> | Contains mutation in *fliC* resulting in the inability to form flagella. | LB supplemented with 100 µg mL<sup>-1</sup> ampicillin at 37°C | Strain is from the Keio collection (42), which was provided by Dr. Robert Landick and the plasmid was provided by Dr. Rodney Welch (43) |
| *E. coli* K-12-Δ*fliC*-46 | BW25113 | GFP gene expression plasmid pGEN-GFP(LVA); Amp<sup>r</sup> | Contains mutation in *fliC* resulting in the inability to form flagella. | LB supplemented with 100 µg mL<sup>-1</sup> ampicillin at 37°C | Strain is from the Keio collection (42), which was provided by Dr. Robert Landick and the plasmid was provided by Dr. Rodney Welch (43) |
| *E. coli* K-12- Δ*fliD*-45 | BW25113 | GFP gene expression plasmid pGEN-GFP(LVA); Amp<sup>r</sup> | Contains mutation in *fliD* resulting in the inability to form flagella. | LB supplemented with 100 µg mL<sup>-1</sup> ampicillin at 37°C | Strain is from the Keio collection (42), which was provided by Dr. Robert Landick and the plasmid was provided by Dr. Rodney Welch (43) |
| *E. coli* K-12-Δ*fliD*-46 | BW25113 | GFP gene expression plasmid pGEN-GFP(LVA); Amp<sup>r</sup> | Contains mutation in *fliD* resulting in the inability to form flagella. | LB supplemented with 100 μg mL<sup>-1</sup> ampicillin at 37°C | Strain is from the Keio collection (42), which was provided by Dr. Robert Landick and the plasmid was provided by Dr. Rodney Welch (43) |
| *E. coli* K-12-Δ*cheB*-3 | BW25113 | GFP gene expression plasmid pGEN-GFP(LVA); Amp<sup>r</sup> | Contains mutation in *cheB* resulting in the inability to chemotaxis. | LB supplemented with 100 μg mL<sup>-1</sup> ampicillin at 37°C | Strain is from the Keio collection (42), which was provided by Dr. Robert Landick and the plasmid was provided by Dr. Rodney Welch (43) |
| *E. coli* K-12-Δ*cheB*-4 | BW25113 | GFP gene expression plasmid pGEN-GFP(LVA); Amp<sup>r</sup> | Contains mutation in *cheB* resulting in the inability to chemotaxis. | LB supplemented with 100 μg mL<sup>-1</sup> ampicillin at 37°C | Strain is from the Keio collection (42), which was provided by Dr. Robert Landick and the plasmid was provided by Dr. Rodney Welch (43) |
| *E. coli* K-12-Δ*cheZ*-59 | BW25113 | GFP gene expression plasmid pGEN-GFP(LVA); Amp<sup>r</sup> | Contains mutation in *cheZ* resulting in the inability to chemotaxis. | LB supplemented with 100 μg mL<sup>-1</sup> ampicillin at 37°C | Strain is from the Keio collection (42), which was provided by Dr. Robert Landick and the plasmid was provided by Dr. Rodney Welch (43) |
| *E. coli* K-12-Δ*cheZ*-60 | BW25113 | GFP gene expression plasmid pGEN-GFP(LVA); Amp<sup>r</sup> | Contains mutation in *cheZ* resulting in the inability to chemotaxis. | LB supplemented with 100 μg mL<sup>-1</sup> ampicillin at 37°C | Strain is from the Keio collection (42), which was provided by Dr. Robert Landick and the plasmid was provided by Dr. Rodney Welch (43) |

<sup>a</sup> Abbreviations for antibiotic resistance are as follows: Em<sup>r</sup>-erythromycin resistance, Cm<sup>r</sup>-chloramphenicol resistance, Amp<sup>r</sup>-ampicillin resistance, Kan<sup>r</sup>-kanamycin resistance, Sp<sup>r</sup>-spectinomycin resistance, Sm<sup>r</sup>-Streptomycin resistance.

### Bacterial sample preparation and imaging

We centrifuged bacterial cells from overnight cultures, washed the cell pellets twice with 1X PBS, and resuspended them in 0.1X tryptic soy broth (TSB). We inoculated *F. johnsoniae* at a cell density of 10^7^ CFU/mL or optical density (OD_600_) = 0.008, with or without other bacterial species. For co-zorbing experiments, we co-inoculated 10^7^ CFU/mL of *F. johnsoniae* with the following bacterial species at these respective cell densities: *E. coli* (10^7^ CFU/mL or OD_600_ = 0.125), *S. aureus* (10^7^ CFU/mL or OD_600_ = 0.1), *L. monocytogenes* (10^7^ CFU/mL or OD_600_ = 0.016), *S. enterica* (10^7^ CFU/mL or OD_600_ = 0.04), *A. tumefaciens* (10^4^ CFU/mL or OD_600_ = 0.0001), *B. cereus* (10^4^ CFU/mL or OD_600_ = 0.000167), *B. subtilis* (10^4^ CFU/mL or OD_600_ = 0.0002), and *P. aeruginosa* (10^6^ CFU/mL or OD_600_ = 0.0049). For tri-zorbing experiments, we co-inoculated 10^7^ CFU/mL of *F. johnsoniae* with 10^7^ CFU/mL of *E. coli* and 10^6^ CFU/mL or OD_600_ = 0.0049 of *P. aeruginosa*. We mixed 1:500 dilution of fluorescently labeled 2 µm diameter polystyrene beads (Bangs Laboratories Cat# FSPP005) with our bacterial samples to test the specificity of co-zorbs. We seeded 2 µL of each sample onto an under-oil microfluidic device (described below) and imaged over 17 hours at room temperature. For staggered inoculation experiments, we seeded 2 µL of 10^7^ CFU/mL of *F. johnsoniae* in the device, allowing zorbs to form for approximately 10 hours. Then, we added 0.2 µL of 10^8^ CFU/mL of *E. coli* or *S. aureus* to the pre-formed zorbs and imaged the samples for 3 hours on a Nikon TI 449 Eclipse inverted microscope. Similarly, we added 0.2 µL of 10^5^ CFU/mL of *E. coli* or *S. aureus* to the pre-formed zorbs and imaged the samples at higher magnification (30x) for ∼30 min at 15 s-intervals using a Zeiss Axio Observer microscope running Zeiss Zen Pro 3.6 software.

Two-dimensional, in vitro time lapse imaging was performed on one of two fluorescence microscopes. The large majority (all that did not require visualization of *P. aeruginosa* and thus a cyan channel) was performed on a Nikon TI 449 Eclipse inverted microscope equipped with brightfield, red and green channels. In experiments requiring visualization of *P. aeruginosa* and thus a cyan channel, imaging was performed on a Zeiss Axio Observer fluorescence microscope. Three-dimensional reconstruction of in vitro zorb/co-zorb structures was performed on z-stacks (0.3-µm step size) obtained on a Nikon AR1 laser scanning confocal microscope courtesy of the UW Optical Imaging Core. Images were obtained for five different co-zorbs and a representative example is shown.

### Preparation of the under-oil open microfluidic system (UOMS) device

Glass chambered cover glass slides were grafted with PDMS-silane through chemical vapor deposition to generate a hydrophobic surface as previously described (9-11). Slides were overlaid with a PDMS stamp containing circular patterns of 2-mm diameter and exposed to O_2_ plasma to etch away the hydrophobic PDMS-silane coating in defined regions (circular spots 2-mm diameter). The now-patterned chambered cover glass slides were overlaid with fluorinated oil (40 cSt) and 2 uL of bacterial cultures were seeded onto the circular spots.

### In vitro image processing and data analysis

Three-dimensional reconstruction of co-zorbs was performed in open-source ICY software. Tracking of zorb and co-zorb motility in vitro was performed on 5-hour timelapse movies (dt = 30 min) using the TrackMate plugin in FIJI/ImageJ (44) using the Laplacian of Gaussian object detector following pre-processing with a median filter, with estimated object diameters ranging from 40 to 100 µm, depending on each movie/species. Tracks were obtained using the Linear Assignment Problem (LAP) algorithm to allow capture of merging events. Tracks were filtered by length for each individual video per the built-in TrackMate algorithm and exported as .csv files containing track position (x,y) and track ID. Mean speeds were calculated for each track using custom MATLAB code. Plots of track speed contain the mean speed for an individual movie (the mean of all individual track means). Plots of track displacement were generated using custom MATLAB code to transform the origin of each track to the origin of the plot (x = y = 0).

Core area fraction and co-zorb circularity (4pi*area/perimeter^2) were calculated in ImageJ by manually outlining the outer perimeter of co-zorbs and their corresponding cores (n = 10 co-zorbs for each movie). Core centrality was calculated as the distance between the core center of mass and the surrounding zorb center of mass; this value was then divided by the mean radius (calculated as the square root of area over pi) of the zorb to yield a measure ranging between 0 (core is directly within the center of the zorb) and 1 (center of the core is located along the outermost edge of the zorb). These measurements were performed on 10 zorb/co-zorbs per movie, with movies from three independent biological replicates for each species.

### Animal ethics statement

Animal care and use protocol M005405-A02 was approved by the Institutional Animal Care and Use Committee (IACUC) at University of Wisconsin-Madison College of Agricultural and Life Sciences (CALS). This protocol adheres to the guidelines established by the federal Health Research Extension Act and the Public Health Service Policy on the Humane Care and Use of Laboratory Animals, overseen by the National Institutes of Health (NIH) Office of Laboratory Animal Welfare (OLAW).

### Zebrafish husbandry and maintenance

Wild-type AB adult zebrafish were maintained under a light/dark cycle of 14 hours and 10 hours, respectively. For experiments, adult fish were in-crossed; embryos were collected and transferred to E3 media (E3) containing Methylene Blue (MB), and kept at 28.5°C. To prevent pigment synthesis and facilitate live-imaging, larvae were switched to E3 -MB + 0.2 mM N-phenylthiourea (PTU, Sigma) starting at 1-day post-fertilization (dpf).

### Bacterial microinjections

Larvae (2dpf) were manually dechorionated and anesthetized in E3 + 0.2 mg/mL Tricaine (ethyl 3-aminobenzoate, Sigma). Using a microinjector, 3 nL of bacterial suspension of *F. johnsoniae, E. coli* and *S. aureus* at cell density of 10^9^ or 10^10^ CFU/mL was injected through the otic vesicle and into the hindbrain ventricle as previously described (16, 17). Bacterial suspensions were mixed with 1% Phenol Red in a 9:1 ratio to facilitate visualization of the inoculum in the hindbrain. After injection, larvae were rinsed three times with E3 -MB) and maintained in E3-MB + PTU for the duration of the experiments.

### In vivo imaging and analysis

Larvae were anesthetized and mounted in agarose (17, 45) in an orientation that allowed full visibility of the hindbrain. To track zorb motility in vivo, z-stacks (3.45 µm step size) were obtained every 6 min on a spinning disk confocal microscope; either a CSU-X; Yokogawa on a Zeiss Observer 399 Z.1 inverted microscope and an electron-multiplying charge-coupled device Evolve 512 camera 400 (Photometrics); or a Nikon A1R 451 inverted Ti2 microscope courtesy of the UW-Madison Optical Core. Three-dimensional reconstructions were generated using ICY software (as above) on z-stacks of step size 0.6 µm. Tracking was performed using the manual tracking plugin within FIJI/ImageJ, and track velocities extrapolated from x,y position data using custom MATLAB code (as above).

## Acknowledgments

This study was supported by the US Army Research Laboratory and the US Army Research Office under Contract/Grant W911NF1910269. J.S. was supported by National Institutes of Health grant T32GM135066-05. N.M.S was supported by National Science Foundation Graduate Research Fellowship Program grant DGE-2137424. J.S, C.L, D.J.B were supported by National Institutes of Health grant NIH P30CA014520. A.H was supported by National Institutes of Health grant R35GM118027.

We thank Dr. E.D. Walker, Dr. Mark McBride, Dr. Jason Peters, Dr. J.D. Sauer, Dr. Ju Wang, Dr. Jeri Barak, Dr. Clay Fuqua, Dr. Robert Landick, Dr. Rodney Welch and their labs for providing the necessary strains and plasmids for this study. We thank Julia F. Nepper for her contributions to the initial stages of the project. We thank the UW Optical Imaging Core for the use of their microscopes. We used Biorender and Adobe illustrator to generate all the schematics and illustrations seen in the manuscript, and ImageJ to process all the images.

## Figures and Table

**<u>Movie 1</u> (separate file)**. Tri-zorb formation. Time-lapse video taken at 30 min intervals showing the formation of tri-zorbs over 17 hours. *F. johnsoniae* is labeled red, *E. coli* is labeled green and *P. aeruginosa* is labeled blue. (All channels [here](https://uwmadison.app.box.com/file/1599739274363).)

**<u>Movie 2</u> (separate file)**. *F. johnsoniae* aggregates around *E. coli* and transports it. Time-lapse video taken at 15 s intervals showing the cells of *F. johnsoniae* (bright-field) aggregate around *E. coli* (green), merging with other aggregates and eventually localizing it to the zorb core.

