## Supplementary material for "Co-zorbs: Motile, multispecies biofilms aid transport of diverse bacterial species"

**This PDF file includes:**

Figures S1 to S4
Legends for Movies S1 to S3

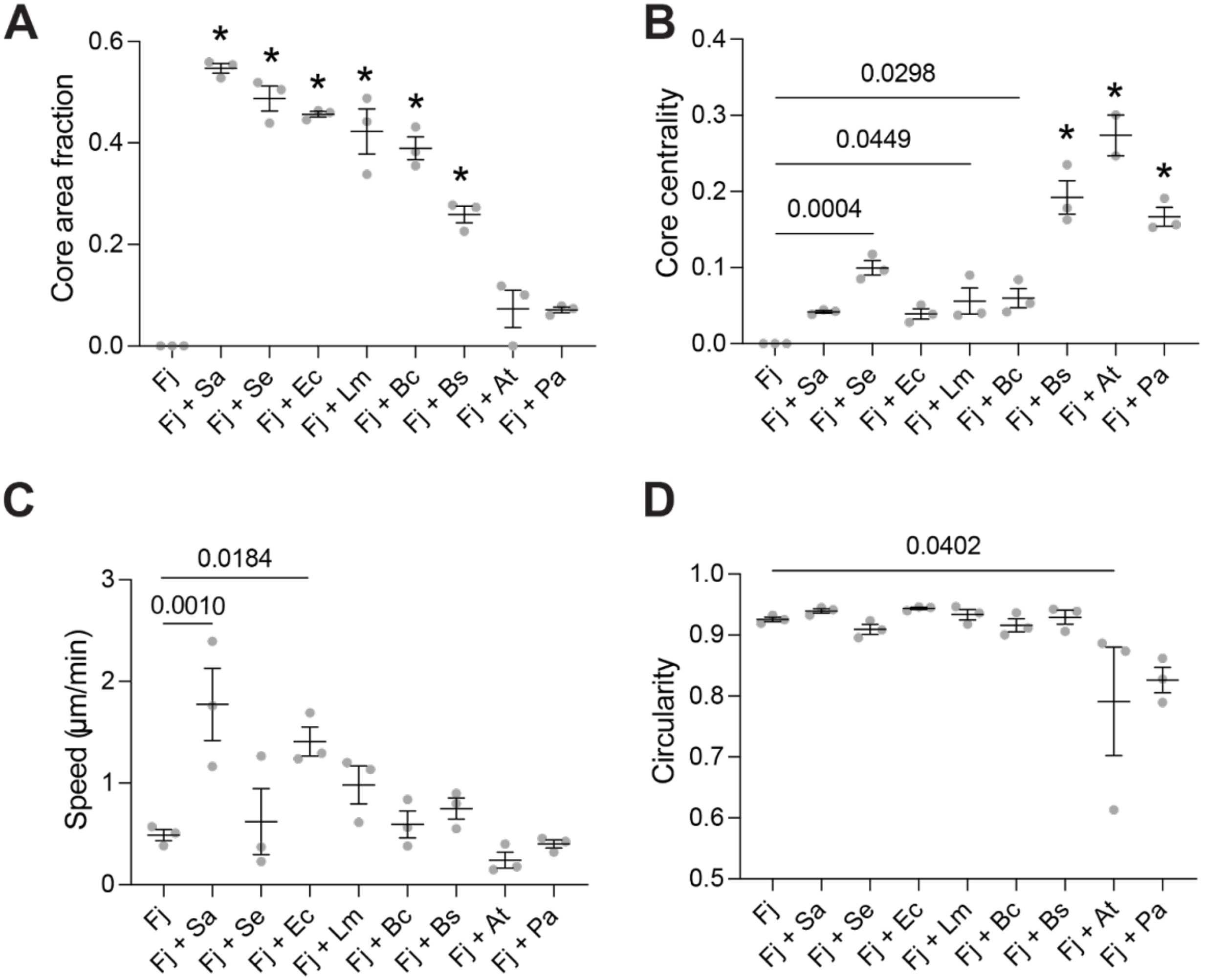


**Fig. S1.** Characteristics of *F. johnsoniae* co-zorbs with various bacterial species. Plots of (A) Co-zorb area fraction (relative to total co-zorb area), (B) Core centrality (distance of core from center normalized by total co-zorb radius) (C) Co-zorb speed and (D) co-zorb circularity, for *F. johnsoniae* co-zorbing with various species. Each point represents the mean of a single biological replicate (n=10 zorbs per biological replicate). Statistical significance was determined by one-way analysis of variance with each condition compared to zorbing (Fj alone). All p-values < .05 are displayed; * denotes p < .0001.

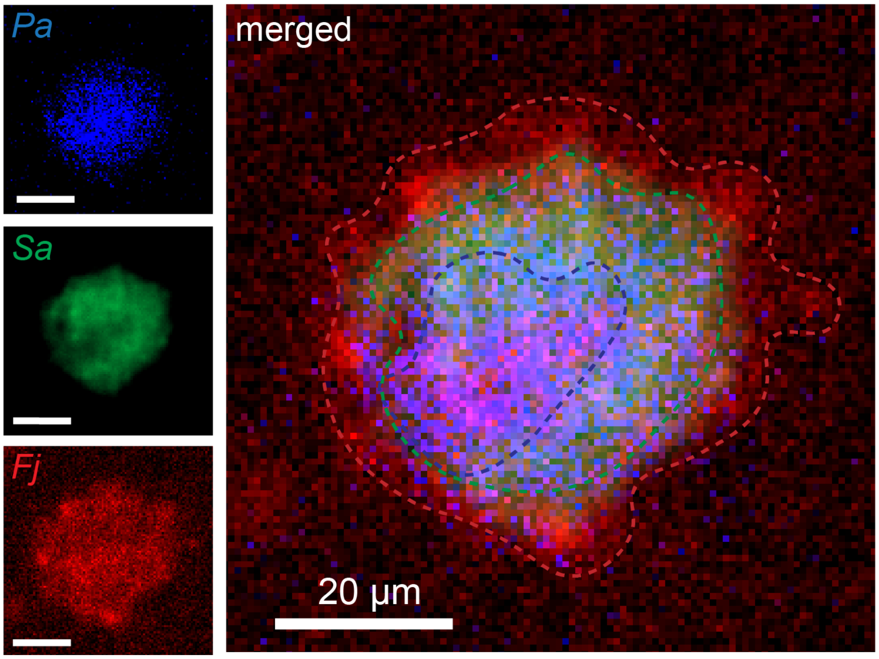


**Fig. S2.** Tri-zorbing with *P. aeruginosa* and *S. aureus*. *F. johnsoniae* is labeled in red (shell), *S. aureus* is in green (outer-core), and *P. aeruginosa* is in blue (inner-core). All scale bars represent 20 µm.

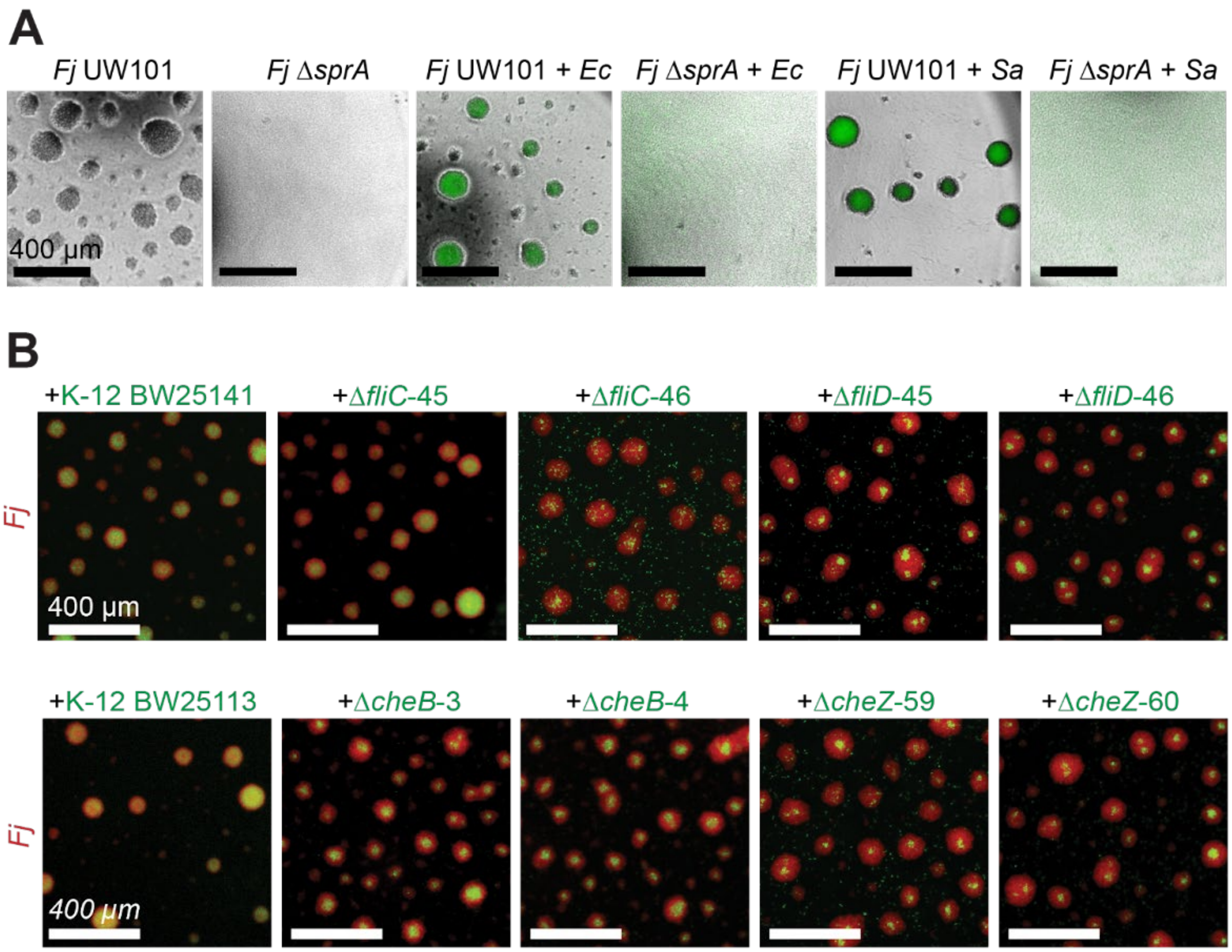


**Fig. S3.** Mutant analysis reveals the role of motility in co-zorbing. (A) Co-zorbing is dependent on motility of *F. johnsoniae*. Images showing zorbs or co-zorbs at ~14 hours showing (left to right) *F. johnsoniae* wild type UW101, Δ*sprA* mutant, *F. johnsoniae*-UW101 with *E.coli*, Δ*sprA* mutant with *E. coli*, *F. johnsoniae*-UW101 with *S. aureus*, and Δ*sprA* mutant with *S. aureus*. (B) Motility of the second species is not necessary for co-zorbing. Top left to right: Co-zorbing between *F. johnsoniae* and *E. coli* wild type K-12 BW25141, Δ*fliC*-45, Δ*fliC*-46, Δ*fliD*-45, Δ*fliD*-46. Bottom left to right: Co-zorbing between *F. johnsoniae* and (i) *E. coli* wild type K-12 BW25113 (ii) Δ*cheB*-3 (iii) Δ*cheB*-4 (iv) Δ*cheZ*-59 (v) Δ*cheZ*-60.

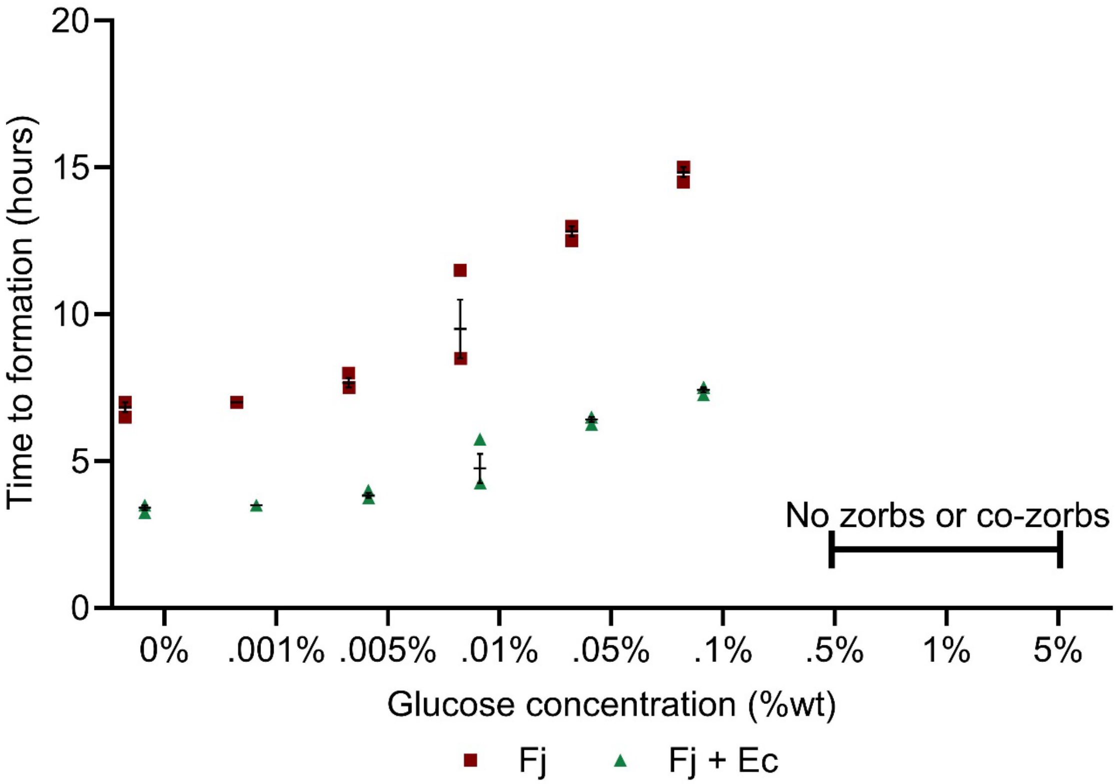


**Fig. S4.** *F. johnsoniae* zorb and co-zorb formation is inhibited in high glucose concentrations. Zorb formation under various glucose concentrations over time for three biological replicates. The error bars denote the mean +/- S.E.M.

**Movie S1 (separate file)**. *F. johnsoniae-E.coli* co-zorb formation. Time-lapse video taken at 30 min intervals showing the formation of co-zorbs over 17 hours. *F. johnsoniae* is labeled red and *E. coli* is labeled green.

**Movie S2 (separate file).** *F. johnsoniae* aggregates around *S. aureus* and transports it. Time-lapse video taken at 15 s intervals showing the cells of *F. johnsoniae* (bright-field) aggregate around *S. aureus* (green), merging with other aggregates and moving it from one point to another.

**Movie S3 (separate file).** Co-zorb formation within zebrafish hindbrain. Z-projection time-lapse videos at 6 min intervals of co-zorbing motility within larval zebrafish (Fj + Ec left; Fj + Sa right), representative co-zorbs are denoted in the first frame by white arrows.
